# Characterization of the Mitomycin C Resistance Protein McrA

**DOI:** 10.1101/2025.11.28.691146

**Authors:** Gwen Tjallinks, Tomás Dunleavy, Tim Jonker, Nicolò Angeleri, Andrea Mattevi, Marco W. Fraaije

## Abstract

McrA from *Streptomyces lavendulae* is a flavin-dependent enzyme thought to provide self-resistance against the DNA-alkylating antibiotic mitomycin C (MMC). McrA belongs to the berberine bridge enzyme (BBE)-like subfamily of oxidases and catalyzes the oxidation of reduced MMC converting it back into the inactive prodrug form, thereby preventing rearrangement into the reactive quinone methide intermediate. Here we demonstrate the first crystal structure of McrA, allowing us to identify key residues involved in retaining MMC in the active site. Biochemical characterization studies such as pre-steady state kinetics verified McrA oxidase activity, while binding studies demonstrated its ability to recognize oxidized MMC.

## Introduction

Mitomycins are a family of natural products produced by streptomycetes.^1^ Members of this family are used as cytotoxic antibiotics because they exhibit strong alkylating activity.^2^ Their activity is connected to their unique chemical skeleton, which is a compact aziridine-containing polycyclic ring decorated with reactive quinone and carbamate moieties. Under reductive conditions and low pH, the mitosane skeleton is prone to activation and can act as a potent DNA alkylating agent (Scheme 1).^3,4^ Mitomycin C (MMC) from *Streptomyces lavendulae* is seen as the most potent alkylating agent and has been utilized in the treatment of various carcinomas including lung, breast and prostate cancer.^5^ A key step in the activation of this prodrug is the two-electron reduction of the quinone moiety of MMC, which triggers a cascade of reactions resulting in the formation of a reactive quinone methide intermediate. This intermediate specifically crosslinks with DNA at 5’-CpG-3’ sequences.

The mitomycin biosynthetic gene cluster from *S. lavendulae* NRRL2564 was discovered by Mao et al.^6^ in 1999 and subsequent studies have revealed the function of the genes related to its skeleton biosynthesis^7–10^ and mitosane tailoring^11–14^. Even though *S. lavendulae* contains a high percentage of G and C bases in its chromosome and is therefore prone to alkylation by MMC, it displays exceptional resistance against MMC.^15^ Several self-resistance genes have been identified that diminish the toxicity of MMC to the host organism, including Mrd, Mct, MitR and McrA (Scheme 1).^16–22^ The drug-binding protein named Mrd was shown to grant MMC resistance when introduced into *Streptomyces lividans* and *Escherichia coli*.^18^ By functioning as a mitomycin-binding protein, it prevents the reductive activation of MMC and thereby subsequent crosslinking of DNA. Presumably, Mrd underwent an evolutionary switch from being a drug-activating quinone reductase to a drug-binding component in the MMC export system.^16^ However, using Mrd as singular cellular self-protection mechanism was thought to be incomplete since most of the MMC was found in the culture medium. *Streptomyces* excrete many secondary metabolites and indeed another MMC resistance determinant was characterized, encoding for a membrane-associated protein named Mct.^19^ Coexpression of Mrd and Mct led to a 150-fold increase in resistance toward MMC, providing evidence that both proteins are part of a mitomycin-specific drug export system. Recently, another self-resistance gene encoding for the berberine bridge enzyme (BBE)-like enzyme MitR was identified.^20^ MitR was found to act as drug-binding and efflux protein, since it contains an N-terminal Tat signal peptide allowing for extracellular protein secretion in *Streptomyces*. Moreover, it was noted that MitR could reduce MMC, which suggests that the protein most likely evolved from a quinone reductase to an MMC binding protein, similar to what was observed for Mrd.

Lastly, the earliest identified self-resistance gene was found in 1994 and named McrA.^21^ McrA is another BBE-like flavoprotein (27% identity to MitR) and was shown to confer MMC resistance by oxidizing the reduced MMC back to its prodrug form (before it can rearrange to the active quinone methide, Scheme 1). As observed for other members of the BBE-like subfamily, McrA contains a covalently-bound FAD cofactor and has a C-terminal BBE domain (pfam entry PF08031).^23,24^ Previous studies observed flavin quenching of McrA when incubated with reduced MMC.^22^ Moreover, it was shown that addition of McrA to reduced MMC prevented DNA crosslinking. In order to have a better understanding of the molecular mechanism that McrA employs, in this study we obtained the first crystal structure of McrA and performed complementary biochemical characterization studies to confirm its role as MMC self-resistance gene. Our functional analysis of McrA, which includes elucidation of its atomic structure, provides insights into the cellular self-defense system of *S. lavendulae* and contributes to a better understanding of McrA and its redox relay mechanism.

**Scheme 1.**
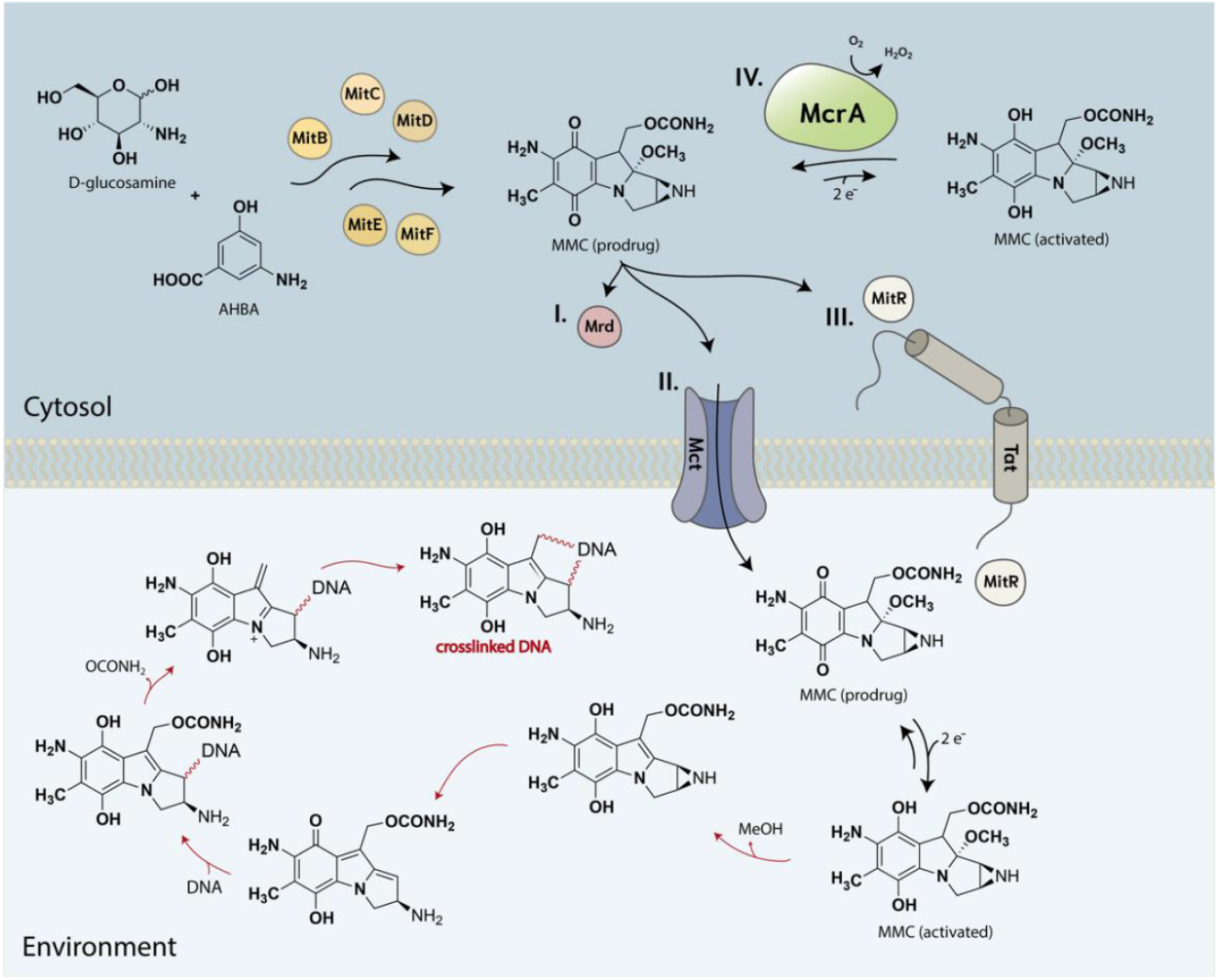
Schematic representation of the proposed cellular self-protection mechanism of *Streptomyces lavendulae* toward mitomycin C (MMC) and consecutive steps for DNA alkylation after MMC activation (red arrows). Self-resistance through utilizing the I. drug-binding protein Mrd. II. membrane-associated protein Mct. III. drug-binding and efflux protein MitR and IV. redox relay mechanism utilized by McrA.

## Results and Discussion

### Biochemical Characterization

McrA was expressed as a soluble protein (63 mg L^-1^) in *Escherichia coli* with a distinct yellow color and typical flavin absorbance spectrum with maxima at 365 nm and 454 nm (Figure 1A and S1). Expression of McrA with covalently-bound FAD was confirmed by acetic acid staining of the SDS-PAGE gel and visualization of UV fluorescence of the flavin cofactor (Figure S2A). The FAD incorporation for McrA was measured and had an Abs_280_/Abs_454_ ratio of 10. To begin with biochemical characterization of McrA, the thermostability of McrA was explored in different buffers, with pH values from 5 to 9, using the thermoFluor method. The highest melting temperature (T_m_) recorded was 59.5 °C in KPi buffer (pH 7). The lowest T_m_ recorded was 33 °C in citric acid buffer (pH 5) (Figure 1B). McrA showed increased thermostability in neutral to more basic buffers, which is a commonly observed phenomenon for intracellular proteins. This pH range is optimal for preserving the enzyme’s tertiary structure and stabilizing amino acid interactions necessary for catalytic activity.

**Figure 1.**
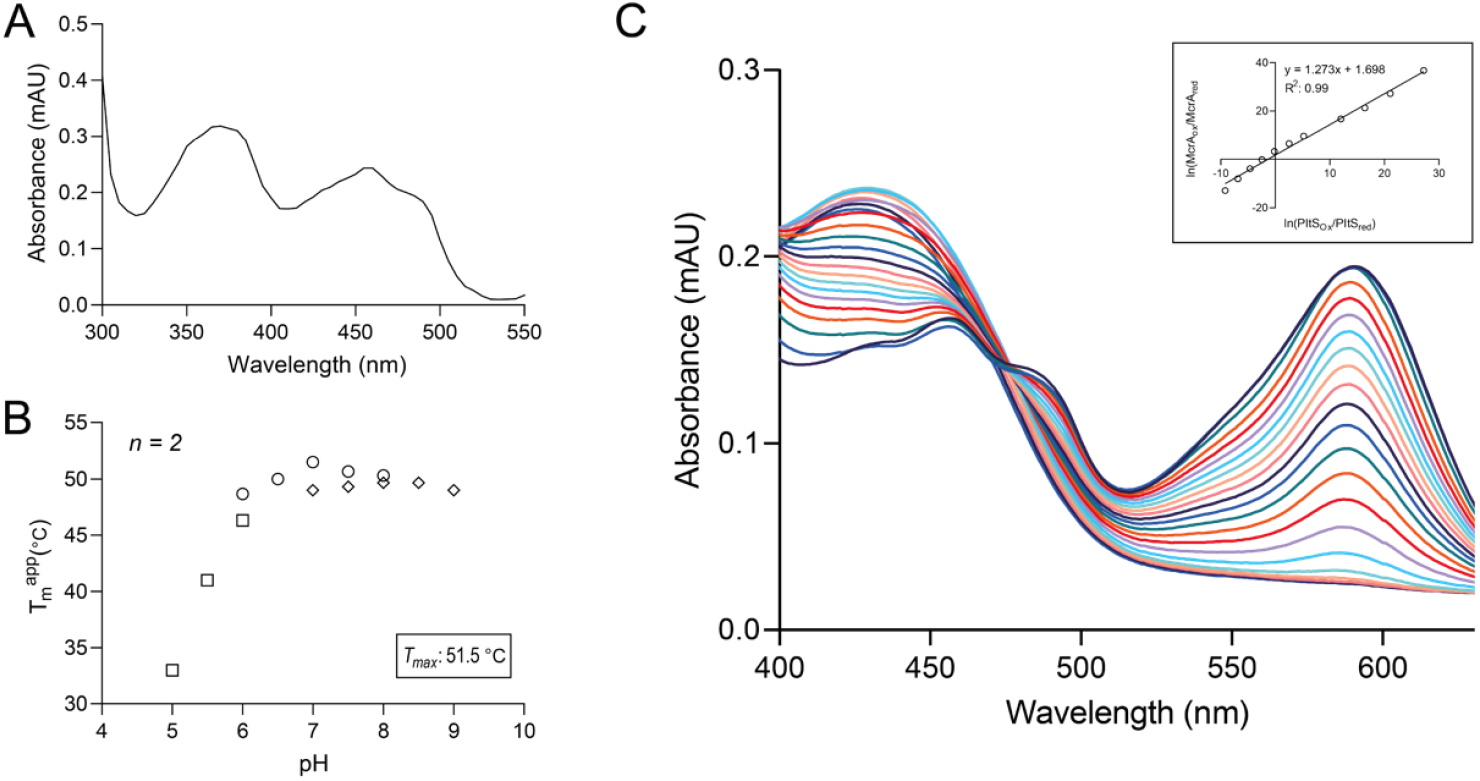
Melting temperatures, FAD absorbance spectrum and redox potential of McrA. (A) 20 μM McrA in 50 mM TRIS-HCl pH 8. (B) Melting temperature profile of McrA at different pH values. Citrate buffer was used for acidic pH (pH 5.0 – 6.0), KPi buffer for neutral pH (pH 6.0 – 8.0), and TRIS-HCl buffer for basic pH (pH 7.0 – 9.5). (C) Determination of the midpoint redox potential by observing the spectral changes of McrA using PITS as dye. They were measured using a scan rate of 1000 nm/min for 60 minutes. The inset displays a plot with the log[(OxMcrA)/(RedMcrA)] on the Y-axis and the log[(OxPITS)/(RedPITS)] on the X-axis. The redox potential of McrA was found to be −35 ± 5 mV.

Using the xanthine/xanthine oxidase system^25^, the midpoint potential (E_m_) of McrA was determined to be −35 ± 5 mV. The reference dye used for this redox titration experiment was indigo tetrasulfonate (PITS), which has an absorbance maximum at 590 nm (Figure 1C). A redox potential of −35 mV is in accordance with the recorded redox potentials of other monocovalent flavoproteins, which typically have E_m_ in the range of −100 mV to +100 mV.^26– 29^ The flavin reduction occurred via a single two-electron transfer process as evidenced from the absorbance spectra recorded during the redox titration. No spectral intermediates were observed that could suggest single electron transfer processes.

To confirm that McrA is indeed an oxidase, as suggested in literature, the reoxidation of the reduced enzyme by molecular oxygen was studied in stopped-flow experiments. McrA was anaerobically reduced via the xanthine/xanthine oxidase system, after which the reduced enzyme was mixed in the stopped-flow instrument with a buffer containing different concentrations of dioxygen. A rapid reoxidation of the flavin cofactor in McrA was observed. The spectra could be fitted well using a first order rate function (a → b, Figure 2A). The obtained kinetic data were plotted and fitted using linear regression, resulting in determined reoxidation rate constant (*k*_ox_) of 1.35 x 10^5^ M^-1^ s^-1^ (Figure 2B). To compare, the well-studied vanillyl-alcohol oxidase (VAO) and glucose oxidase (GOX) have a reoxidation rate of 3.1 x 10^5^ M^-1^ s^-1^ and 1.5 x 10^6^ M^-1^ s^-1^ (at low pH) respectively.^30–32^ Such rapid rate of flavin reoxidation by molecular oxygen confirms that McrA acts as an efficient flavoprotein oxidase. The reoxidation of McrA in the presence of oxidized MMC (270 µM) was also recorded, but no dramatic change in *k*_obs_ was observed (Fig 2B). It is known that reduced MMC is a labile compound that can undergo a spontaneous elimination of methanol, forming a highly reactive electrophilic metabolite.^33^ Therefore, due to the instability of the substrate, it was not possible to monitor the process of flavin reduction.^34^

**Figure 2.**
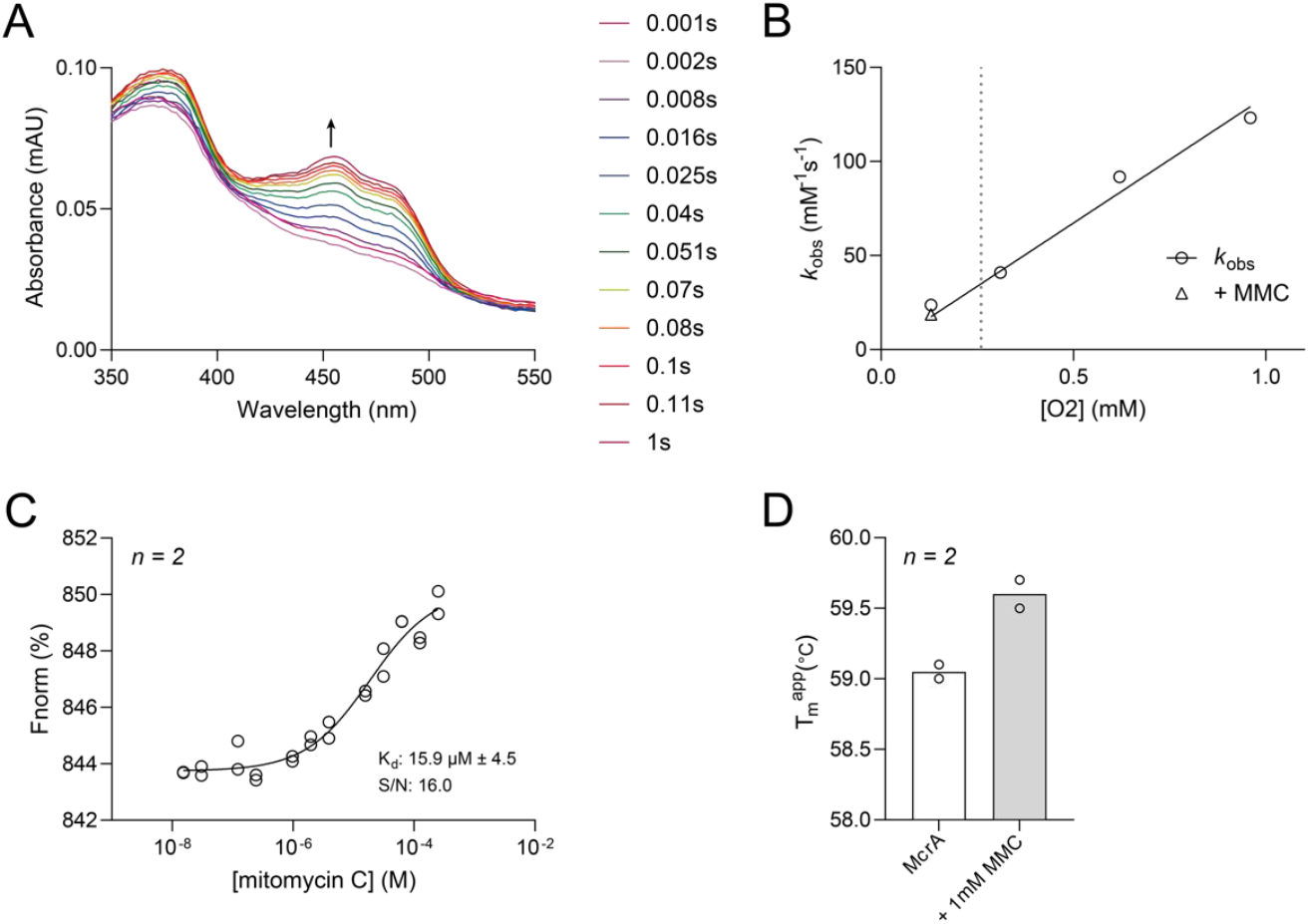
Pre-steady state kinetics and binding affinity of McrA toward mitomycin C (MMC). (A) Spectral changes observed upon aerobic mixing of reduced McrA with buffer containing dioxygen. (B) The oxidative half-reaction of McrA with different dioxygen concentrations and with addition of oxidized MMC (270 µM), *n* = *4*. The dotted line indicates the atmospheric oxygen concentration. (C) The dissociation curve of MMC with McrA. The analysis was performed using microscale thermophoresis. (D) Label-free measurements of thermal shifts induced by 1 mM MMC to McrA using Tycho NT.6.

Binding of MMC by McrA was confirmed with microscale thermophoresis and showed that McrA binds MMC with a *K*_*D*_ of 15.9 ± 4.5 µM (Figure 2C). This indicates relatively tight binding and is within a similar range compared to the MMC-binding proteins MitR (58.52 ± 5.64 μM) and Mrd (53.98 ± 4.58 μM).^20^ Moreover, binding of MMC led to a slight stabilization in melting temperature of McrA by 0.5 °C (Figure 2D). Although the used MMC was in the oxidized form, it is expected that McrA displays a similar or higher affinity towards the reduced activated form of MMC, as the two compounds are highly similar.

### Structural Analysis

McrA exists as a monomer in solution, confirmed by size-exclusion (Figure S2B). McrA crystallized with two monomers in the asymmetric unit, and the general structure was solved at 2.1 Å (Table S1, PDB 9R43). McrA is part of the vanillyl-alcohol oxidase/*p*-cresol methylhydroxylase (VAO/PCMH) family and belongs specifically to the BBE subfamily.^35,36^ The general structure of McrA consists of two main domains: the N-terminal FAD-binding domain (residues 27 – 195) and the C-terminal substrate-binding domain (residues 196 – 409). McrA also contains a short C-terminal BBE domain (residues 410 – 442, Figure 3A).^24^

**Figure 3.**
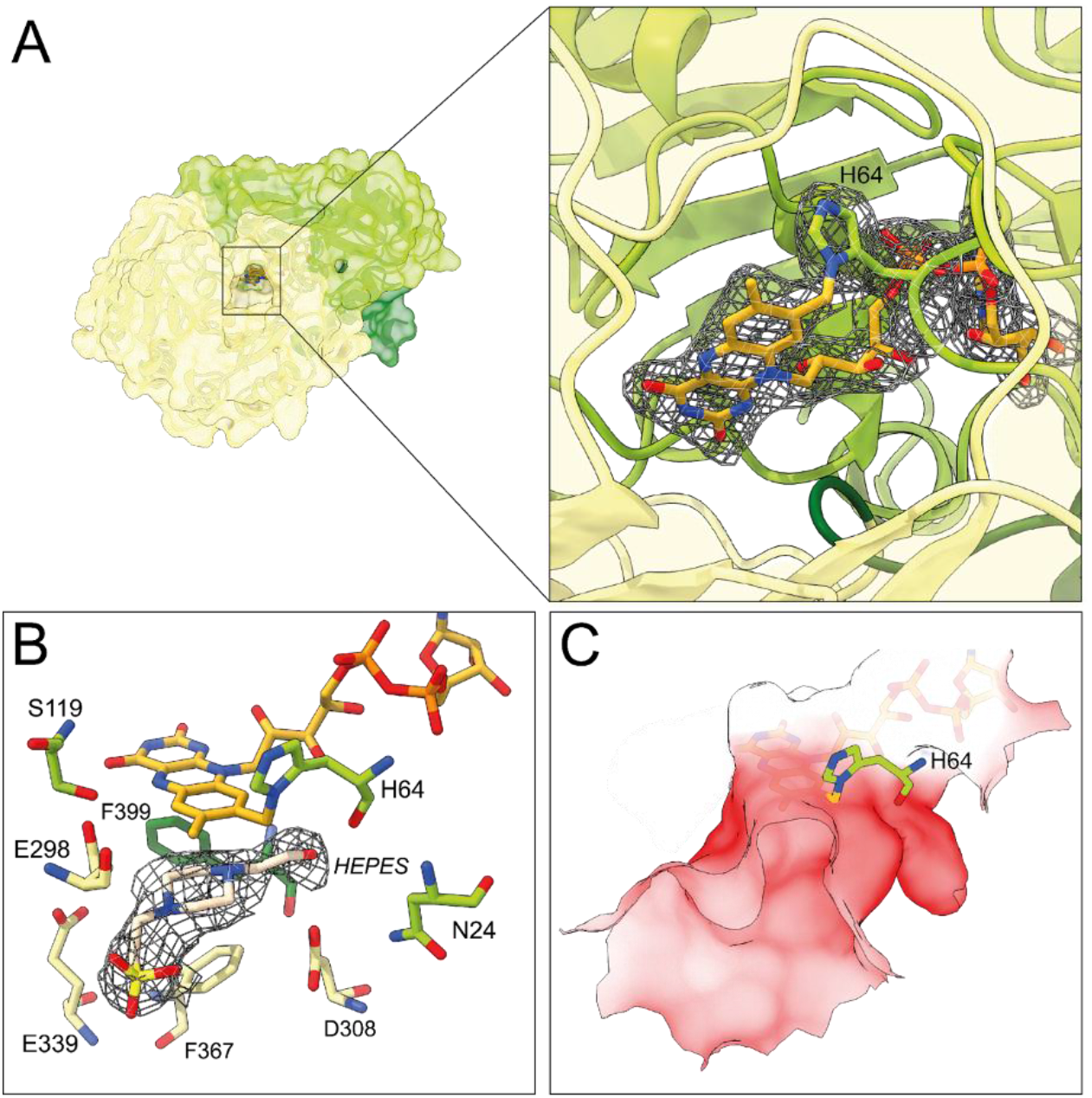
Crystal structure of McrA shown as ribbon diagram (PDB 9R43). (A) Monomeric representation of McrA with the substrate binding domain in light yellow, the FAD binding domain in light green and the BBE domain in dark green. A close-up image of the covalently-bound FAD to His64 is displayed including their electron density. The FAD and His64 polder omit map was calculated by excluding the cofactor and residue and was contoured at 4.5 σ. (B) A HEPES molecule bound in the active site of chain B. The HEPES polder omit map was calculated by excluding the molecule and contouring it at 4.6 σ. (C) The negative electrostatic potential of the active site of McrA, displayed in red.

A common characteristic for BBE-like oxidases is the incorporation of a covalently-bound flavin cofactor.^37^ This feature was also observed in the structure of McrA, where the FAD is covalently anchored to His64 (Figure 3A). Covalent incorporation of FAD allows for a more open active site that can assist with the entry of bulkier substrates and it is reasonable to assume that this also aids in MMC oxidation. Moreover, covalent flavinylation generally also results in a relatively high redox potential (as observed for McrA). We obtained the structure of McrA with acetate bound in the active site in monomer A and HEPES bound in monomer B (Figure 3B), which were part of the crystallization solution. Moreover, HEPES buffer (50 mM, 30 mM NaCl, pH 7) was used for size-exclusion chromatography as sample preparation for crystallization plates. Native MS indicated that HEPES is most likely binding to the surface of McrA, since multiple peaks were found with a mass difference corresponding to the mass of HEPES. As control, size-exclusion was performed using KPi buffer (50 mM, pH 7), which resulted in a single peak corresponding to the molecular weight of flavin-bound SUMO-cut McrA (49390 Da, Figure S3).

The active site of McrA is decorated with a multitude of negatively charged amino acids, including Glu298, Asp308 and Glu339 that contribute to its negative electrostatic potential (Figure 3C). This negative electrostatic potential could potentially help retain MMC in the active site. The crystal structures of mitomycin 7-O-methyltransferase (MmcR, PDB 3GXO) and Mrd (PDB 1KLL) showed to have mitomycin bound in a similar negatively-charged pocket. Reductively activated MMC was shown to exhibit strong electrostatic interactions to polyanions, hence these moieties could form non-covalent electrostatic interactions with the negatively charged residues present in the active site.^38,39^ The active site is lined with a combination of hydrophobic and polar (un)charged residues. Hydrophobic and uncharged residues include Ser119, His296, Pro299, Tyr304, Val337, Phe367, Gly369 and Phe399. The negative and positively charged residues are Arg256, Asp308, Glu339, Arg341 and Glu298.

In order to confirm MMC binding, several substrate soaking crystallography trials were performed. Unfortunately, a crystal structure of McrA with (oxidized) MMC bound in the active site could not be obtained, possibly because affinity for the reduced form is much higher. Nevertheless, by molecular docking studies we could identify potentially crucial active site residues interacting with activated MMC (Figure 4). First of all, the mitosane skeleton of MMC was sandwiched between the isoalloxazine ring of FAD and residue Phe367, allowing for placement of the C8 hydroxyl group in close proximity to the N5 atom of the flavin cofactor. The negatively charged Asp308 can form a hydrogen bond with the C5 hydroxyl group of the hydroquinone ring and can therefore act as a potential catalytic base. Moreover, this residue might be involved in forming hydrogen bond interactions with the nitrogen atom of MMC’s aziridine ring. The second negatively-charged residue Glu339 establishes a hydrogen bond with the amine group of MMC’s hydroquinone ring and the third negatively-charged residue Glu298 might provide stabilizing interactions to the C8 hydroxyl group of MMC. Moreover, a positively-charged Arg256 provides hydrogen bonding with the carbamate side-chain of MMC. Several other residues were highlighted that contribute to additional hydrophobic and van der Waals interactions, including Pro299, Val337, Gly369 and Phe399.

**Figure 4.**
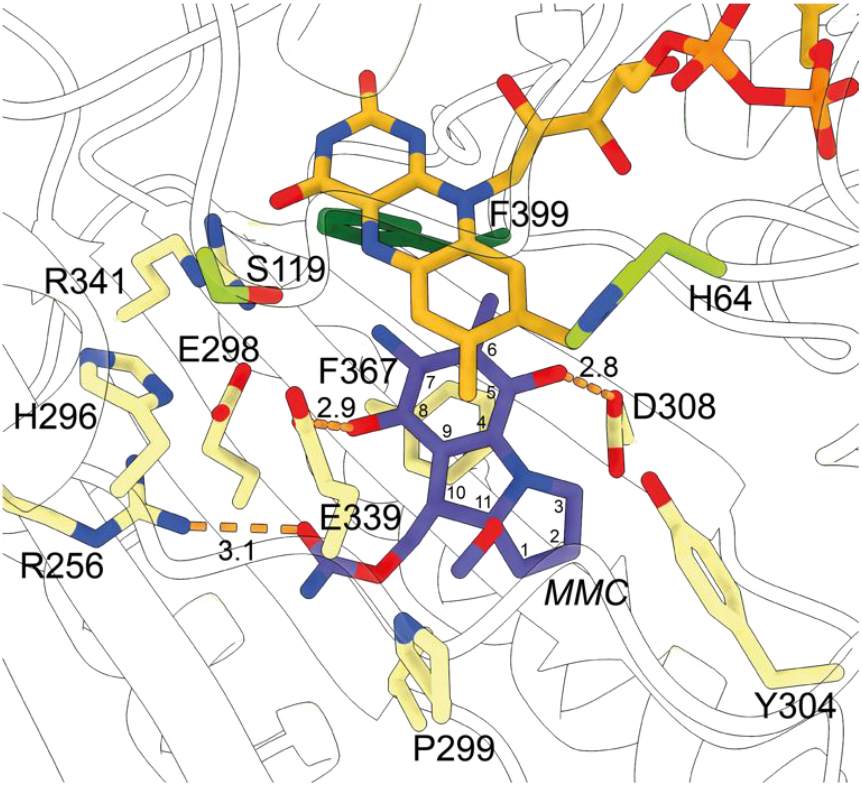
McrA active site entrance and docking studies with MMC. Docking of MMC highlighting hydrogen-bonding interactions with active site residues including Arg256, Asp308 and Glu339. Distances of hydrogen-bonding interactions are shown in Å.

Due to the labile nature of reduced MMC, the oxidase activity of McrA could not be documented. Nevertheless, pre-steady state kinetics demonstrated that the reduced flavin of McrA rapidly reoxidized in the presence of molecular oxygen (1.35 × 10^5^ M^-1^ s^-1^). Moreover, our work on McrA has shown that this flavoenzyme is able to bind the oxidized prodrug MMC. By solving the crystal structure of McrA, we could identify a multitude of negatively charged amino acids, including Glu298, Asp308 and Glu339 that contribute to it negative electrostatic potential. Other MMC-binding proteins, including Mrd and MmcR, displayed similar negatively-charged active site pockets and it is thought to help retain MMC. Lastly, docking studies revealed that MMC fits well within the active site of McrA, offering structural insights into how the enzyme mediates MMC oxidation to mitigate autotoxicity.

## Summary

McrA is a flavin-dependent enzyme that protects *Streptomyces lavendulae* from the cytotoxic antibiotic mitomycin C by re-oxidizing its activated, toxic form back into the inactive prodrug. This study reports the first crystal structure of McrA and identifies key residues that help retain MMC in its negatively charged active site. It is also shown that the FAD-containing enzyme is monomeric and able to efficiently use dioxygen as electron acceptor, demonstrating that it acts as an oxidase. Together, the findings clarify McrA’s role in the organism’s self-resistance machinery.

## Materials and Methods

### Chemicals

All commercial chemicals for assays were purchased from Sigma-Aldrich, except for K_2_HPO_4_ and KH_2_PO_4_ that were purchased from Carlo Erba Reagents. Mitomycin C was purchased from Fischer Scientific. NaCl was purchased from VWR Chemicals. Acetonitrile Ultra LC and Water Ultra LC were purchased from Romil.

### Cloning and Transformation

The *Escherichia coli* codon-optimized gene of McrA from *Streptomyces lavendulae* (Genbank ID: AAA21476.1) was ordered from Twist Biosciences containing BsaI sites at the 5’ and 3’ termini. Cloning of the gene into a pBAD vector containing an N-terminus His_6_-SUMO tag was performed using the Golden Gate methodology^40^ with a T100 Thermal Cycler (Bio-Rad). 100 ng PCR product was added to 50 µL *E. coli* NEB® 10-beta rubidium chloride competent cells and left on ice for 30 min. Heat shock was performed at 42 °C for 45 sec after which the cells were immediately placed on ice for 5 min. Next, 500 µL LB was added and the cells were incubated at 37 °C and 200 rpm for 45 min. Then, cells were plated on LB agar plates containing 50 µg mL^-1^ ampicillin and left at 37 °C overnight. Colonies were chosen and grown in 5 mL LB supplemented with 50 µg mL^-1^ ampicillin at 37 °C overnight. Plasmid isolation was performed using the QIAprep® Spin Miniprep Kit (QIAGEN) and Sanger sequencing (Eurofins) was done in order to confirm successful cloning.

### Expression and Purification

Expression of McrA was accomplished by growing a 5 mL LB culture supplemented with 50 µg mL^-1^ ampicillin at 37 °C overnight. Next, the culture was resuspended in 500 mL terrific broth (TB) medium containing 50 µg mL^-1^ ampicillin and shaken at 37 °C and 200 rpm until an OD_600_ between 0.6 – 0.8. Then, induction was initiated with 0.02% L-arabinose and incubated overnight at 24 °C and 200 rpm. Cells were harvested by centrifugation at 5000 rpm, 4 °C for 15 min (Eppendorf 5920R), resuspended in lysis buffer (50 mM KPi, 500 mM NaCl, 5% w/w glycerol, pH 7.5) supplemented with 0.1 µM phenylmethylsulfonyl fluoride, and disrupted by sonication (3 s on, 5 s off, 70% amplitude, 10 min; Sonics Vibracell). Cell debris was removed by centrifugation at 12,000 rpm, 4 °C for 55 min (Eppendorf 5920R). A Ni-Sepharose column (Cytiva) was equilibrated with lysis buffer and then loaded with supernatant. The column was washed with 3 column volumes (CV) wash buffer (lysis buffer + 20 mM imidazole, pH 7.5). Elution was accomplished by adding 1.5 CV elution buffer (lysis buffer + 500 mM imidazole, pH 7.5). Then, a PD10 desalting column (Cytiva) was used for buffer exchange to storage buffer (50 mM KPi, 150 mM NaCl, 5% w/w glycerol, pH 7.5). SDS-PAGE gel electrophoresis (Bio-Rad) was performed to confirm purity and showed a band around 60 kDa that corresponds to the theoretical molecular weight of His_6_-SUMO-fused McrA. Purification yielded 63.2 mg L^-1^ of McrA, exhibiting a characteristic yellow color and classical flavin absorbance spectrum.

### Thermostability Analysis

The melting temperature of McrA was determined using the ThermoFluor assay. An enzyme stock of 1 mg ml^-1^ was prepared, alongside a 1:100 SYPRO Orange solution. For every pH measurement, 5 µL enzyme stock was added to 15 µL buffer, alongside 5 µL of the SYPRO Orange solution. The relative fluorescence change was then monitored over a temperature range from 20 °C to 99 °C with a gradient increase of 1 °C in a CFX96 Touch Real-Time PCR Detection System (Bio-Rad Laboratories). The same protocol was used to determine the tolerance of McrA towards DMSO as a cosolvent.

The melting temperature of McrA with and without addition of 1 mM mitomycin C (MMC) was performed in duplicate using the TychoTMNT.6 system (NanoTemper GmbH). Each vial contained 20 µL of 20 µM McrA in TRIS-HCl buffer (50 mM, pH 8), with one vial containing an additional 1 mM of MMC. The intrinsic fluorescence of tryptophan and tyrosine residues (emission at 330 nm and 350 nm) was measured to determine the melting curves using a temperature gradient from 35 °C to 95 °C with a 30 °C increment per minute. The inflection temperature (T_m_) was derived from the F_350_/F_330_ ratio.

### Redox Potential

To determine the redox potential of McrA, the xanthine/xanthine oxidase system was employed.^25^ For this, the dye potassium indigo tetrasulfonate (E^O^ = −46 mV, 10 µM) was selected, with a maximum absorbance at 590 nm. 5 µM enzyme was used in KPi buffer (50 mM, pH 7.5). The assay was performed on a Jasco V-660 spectrophotometer, with spectra recorded every 90 s for 30 min. The redox potential od McrA was then calculated using the Nerst equation. All experiments were performed in triplicate.

### Pre-steady state kinetics

To investigate whether McrA acted as a true oxidase, pre-steady state kinetics were measured using a SX20 stopped-flow spectrometer, equipped with a photodiode array (PDA) detector (Applied Photophysics, Surrey, England). Sealed vials were made anaerobic using glucose (10 mM) and glucose oxidase (*Aspergillus niger* Type VII, Sigma Aldrich, cat. equivalent) in KPi buffer (50 mM, pH 7.5), while flushing with nitrogen. Once the system was flushed three times with anaerobic buffer, the flavin cofactor of McrA was reduced using the xanthine/xanthine oxidase system. The flavin reduction was followed by eye, with the yellow enzyme solution turning colorless. The reduced enzyme was then mixed with buffer containing different concentrations of oxygen, and spectra were recorded. The kinetic data obtained was then analyzed using a linear regression, with *k*_ox_ being calculated as 134.5 mM^-1^ s^-1^. All measurements were performed in triplicate.

### Native Mass Spectrometry

To determine the chemical compound that contributed to the electron density found in the active site of the crystal structure of McrA, we employed native mass spectrometry. First, McrA was purified by SEC using two different buffers, namely KPi (50 mM, pH 7) and HEPES (50 mM, pH 7, 30 mM NaCl). Both protein samples underwent buffer exchange using 10 mM ammonium acetate pH 7.2 with a PD Spintrap G-25 (Cytiva). A final protein concentration of 10 µM was prepared in 500 µL 10 mM ammonium acetate pH 7.2. Thereafter, the QTOF mass spectrometer (AB Sciex X500B) equipped with a Turbo V Ion source and Twin Sprayer electrospray ionization (ESI) probe was used to discover whether a difference in mass could be detected for the sample that was gel-filtered using HEPES buffer. For the mass spectra, a positive polarity was set using the parameters listed below. Parameters: 35 psi curtain gas, 300 °C temperature, 100 V declustering potential, 50 psi ion source gas 1, 50 psi ion source gas 2, 4500 V ion spray voltage, 5 V collision energy, TOF mass range 2500 – 5500 Da, accumulation time 0.1 s. The injection speed was set to 100 uL min^-1^. For mass deconvolution, the Bio Tool Kit extension of SciexOS 2.1 was used.

### Protein Crystallization, Structure Determination, and Analysis

For protein crystallization, the His_6_-SUMO tag of McrA was cleaved overnight at 4 °C using 1 mg mL^-1^ His_6_-tagged SUMO protease per 100 mg mL^-1^ of McrA in HEPES buffer (50 mM, pH 7) supplemented with 30 mM NaCl. To purify the cleaved McrA, a reverse His-tag purification was performed. A Ni-Sepharose column (Cytiva) was equilibrated with HEPES buffer (50 mM, pH 7, 30 mM NaCl) containing 30 mM NaCl and the flow-through containing the tag-less McrA was collected. Thereafter, further purification was performed using a Superdex200 100/30 column (Cytiva) equilibrated with HEPES buffer (50 mM, pH 7, 30 mM NaCl) on an Äkta Pure (Cytiva) detecting 280 nm, 365 nm and 450 nm wavelengths. The fractions with a yellow color were pooled and concentrated to 12 mg mL^-1^ with the Amicon Ultra (Merck) 30 kDa cut-off. An Oryx Vizier robot (Douglas Instruments) was employed for the crystallization screening using the sitting-drop method in MCR2 96-well plates (Swissci). Dark-yellow crystals were obtained in the JCSG Core Suite IV screening containing 0.2 M calcium acetate, 0.1 M HEPES pH 7.5 and 40% v/v PEG 400 (Figure S4). The crystals were flash-cooled in liquid nitrogen. The X-ray diffraction data was recorded on the MASSIF-1 beamline at the European Synchrotron Radiation Facility (ESRF, DOI 10.15151/ESRF-ES-1934156944).

At the ESRF the automatic data processing was performed using XIA2_DIALS.^41^ *Phaser*^42^ with AlphaFold3 was used to determine the structure of McrA. Two monomers were found in the asymmetric unit and refinement was performed using *Coot*^43^ and *REFMAC*5^44^. The polder omit map was generated to exclude bulk solvent around HEPES and FAD using Phenix.^45^ Validation of the model was done using the wwPDB Validation Service. UCSF ChimeraX was used for figure preparation.^46^ Details regarding the refinement and data collection statistics are available in Table S1. The structure factor amplitudes and data collection statistics were uploaded in the Protein Data Bank (PDB) with the following PDB ID: 9R43.

### Microscale Thermophoresis

His_6_-SUMO-tagged McrA was labelled using the 2^nd^ generation RED-TRIS-NTA dye (NanoTemper GmbH) following the instructions of the kit. In brief, 400 nM of protein was diluted in a PBS buffer supplemented with 0.05% v/v Tween-20 (PBS-T) and incubated with 50 nM dye for 30 min at RT. In the binding assay, a final protein concentration of 100 nM was used and MMC was titrated in a serial dilution with the concentration ranging from 250 µM to 15 nM using PBS-T as buffer.

### Docking Studies

Molecular docking was performed using AutoDock Vina (version 1.1.2).^47^ The crystal structure of McrA (2.1 Å, PDB 9R43) was used to dock MMC inside the active site pocket. A cube cell size (x = 44, y = 40, z = 8) with a search space volume of > 27.000 Å^3^ was placed around the active site of McrA (x = 46.97, y = −20.46, z = 8.26). Vina docking parameters included an energy range of 4 and an exhaustiveness of 8. UCSF ChimeraX was used for preparing the figures.^46^

## Supporting information

supporting information McrA

## Abbreviations

McrA: mitomycin self-resistance protein
MMC: mitomycin C
BBE: berberine-bridge enzyme
FAD: flavin adenine dinucleotide
VAO/PCMH: vanillyl-alcohol oxidase/*p*-cresol methylhydroxylase

## Author Contributions

G.T. and N.A. performed the crystallization, native mass spectrometry and ligand binding studies. T.D and T.J. performed the pre-steady state kinetics, redox potential determination and thermostability analyses. G.T. and T.D. wrote the paper. A.M. and M.W.F. assisted with writing the paper. All authors reviewed the paper.

## Acknowledgement

We are grateful to A. Gottinger for help with performing the native mass spectrometry experiment and to X. Fan for initial protein purification studies. We thank S. Rovida for assistance with protein crystallization experiments. We acknowledge Centro Grandi Strumenti, University of Pavia for providing the analytical machinery. We are grateful to the ESRF for provision of the X-ray beamtime. This research was supported by the COST Action COZYME (CA21162) from the European Union’s Cooperation in Science and Technology-funded group networking.

## Data Availability Statement

Structural data is openly available in the wwPDB under the accession number 9R43. Additional information supporting the results of this manuscript can be found in the Supplementary Information.

