## supporting information McrA for "Characterization of the Mitomycin C Resistance Protein McrA"

### Table of Contents

**Supporting Tables** .......... 2

**Supporting Figures** .......... 3

### Supporting Tables

**Table S1**. Crystallographic data collection and refinement statistics.

| | McrA |
|---|---|
| PDB entry | 9R43 |
| Wavelength (Å) | 0.96546 |
| Resolution range (Å) | 72.24 - 2.10 (2.16 - 2.10) |
| Space group | $P222_1$ |
| Unit cell (Å) | 77.87 80.49 163.81 |
| Total reflections | 291510 (22436) |
| Unique reflections | 59607 (4643) |
| Multiplicity | 4.9 (4.8) |
| Completeness (%) | 98.4 (99.3) |
| Mean I/sigma(I) | 6.5 (1.41) |
| $CC_{1/2}$ | 0.973 (0.673) |
| $R_{merge}$ | 0.11 |
| $R_{work}/R_{free}$ | 0.202 (0.308)/ 0.247 (0.338) |
| No. of atoms | 7166 |
| - protein | 6643 |
| - ligands | 280 |
| - water | 243 |
| RMS bond length (Å) | 0.0085 |
| RMS bond angles (°) | 1.8304 |
| Ramachandran favored (%) | 93.74 |
| Ramachandran allowed (%) | 5.46 |
| Ramachandran outliers (%) | 0.80 |
| Average B-factor (Å$^2$) | 48.0 |
| - protein | 47.47 |
| - ligands | 53.93 |
| - water | 50.35 |

In parentheses the statistics for the highest-resolution shell are given.

### Supporting Figures

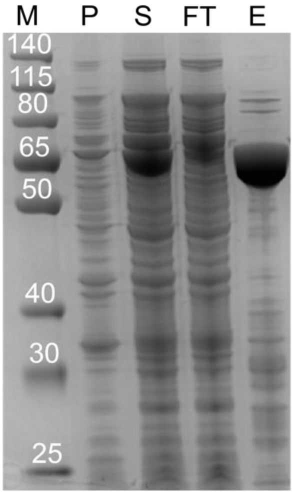


**Figure S1**. SDS-PAGE of purified His$_6$-SUMO-tagged McrA (M: marker, P: pellet, S: supernatant, FT: flow-through, E: elution). The gel was stained with Coomassie Blue R-250.

A

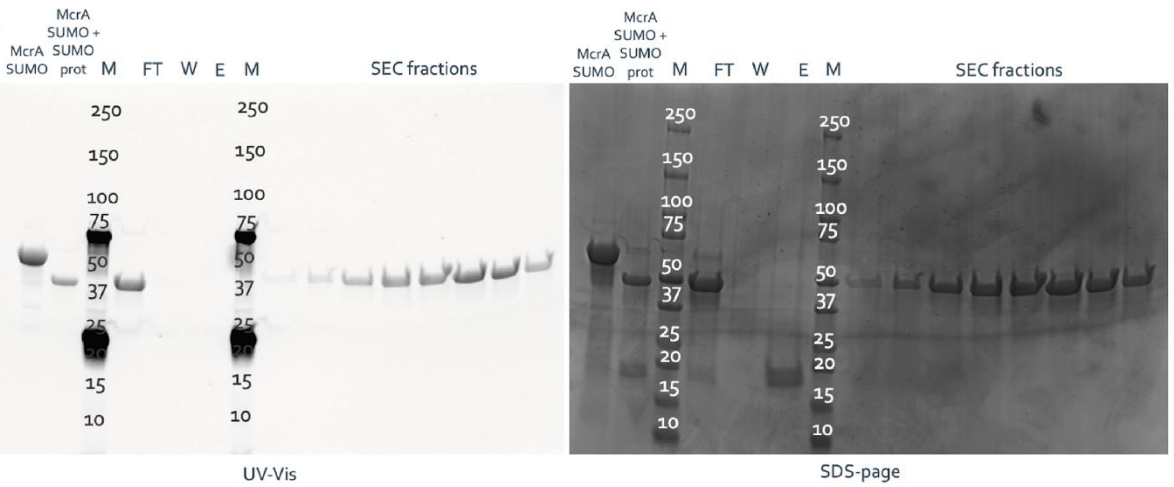


B

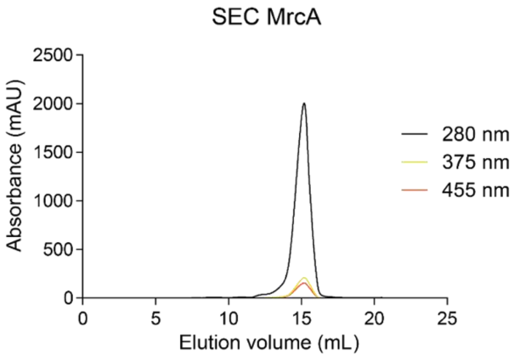


**Figure S2.** Uncropped SDS-PAGE gels of purified His$_6$-SUMO-tagged McrA before and after addition of SUMO-protease and size-exclusion chromatography. (A) Uncropped SDS-PAGE gel of SorC before and after addition of SUMO protease, and different fractions after size-exclusion chromatography (TF: total fraction, W: wash, E: elution). The left gel was stained with acetic acid and visualized by UV-illuminated light of the covalently-bound FAD to His64 of McrA. The right gel was stained with Coomassie blue R-250. (B) The chromatogram of SUMO-cut McrA using the Superdex200 100/30 increase column.

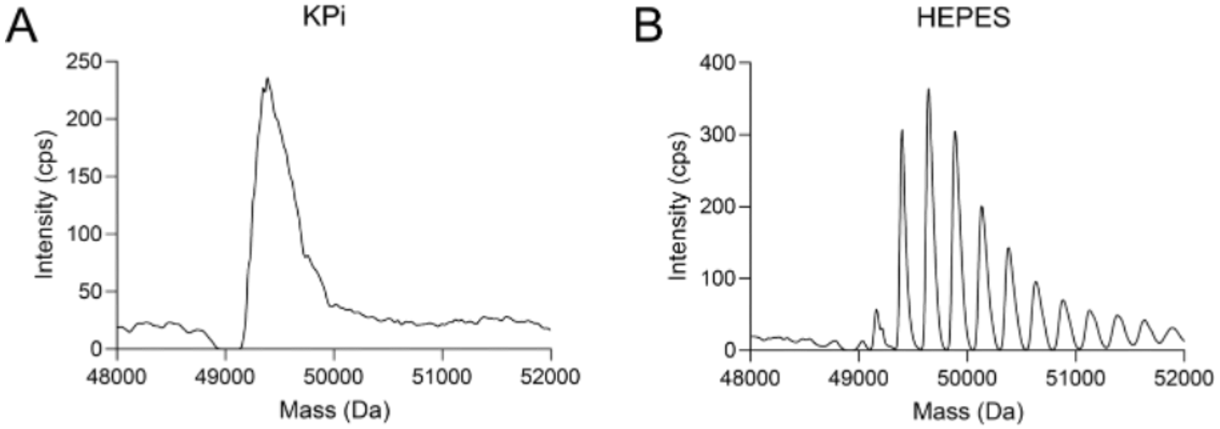


**Figure S3**. Native protein MS. (A) Deconvoluted mass distribution from native protein MS after performing size-exclusion of SUMO-cut McrA in KPi buffer. (B) Deconvoluted mass distribution from native protein MS after performing size-exclusion of SUMO-cut McrA in HEPES buffer. The difference in MW between the peaks roughly corresponds to the molecular weight of HEPES.

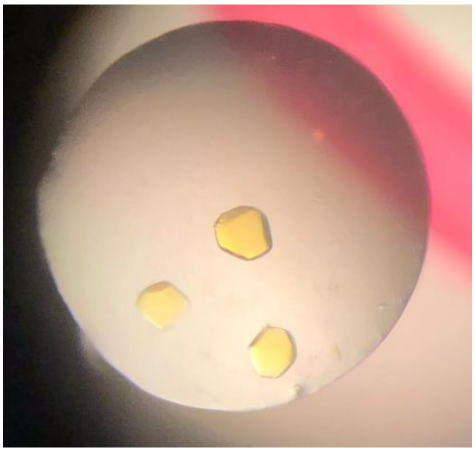

**Figure S4**. Crystals of McrA. Dark-yellow McrA crystals obtained in the JCSG Core Suite IV screening condition containing 0.2 M calcium acetate, 0.1 M HEPES pH 7.5 and 40% (v/v) PEG 400 using 12 mg $mL^{-1}$ McrA.

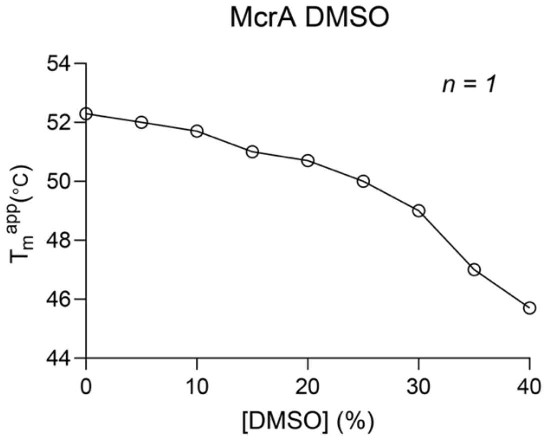


**Figure S5**. Thermostability of McrA with increasing DMSO concentration. A gradual decrease in melting temperature of McrA with increasing v/v% DSMO.
